# Altering a complex serial reaction time task using dual DLPFC and M1-transcranial direct current stimulation

**DOI:** 10.1101/2023.07.26.550620

**Authors:** E. Kaminski, D. Carius, J. Knieke, N. Mizuguchi, P. Ragert

**Author notes:** Correspondence Elisabeth Kaminski, Dr., Department of Movement Neuroscience, Faculty of Sport Science, University of Leipzig, Jahnallee 59, 04109 Leipzig, Germany.

## Abstract

Transcranial direct current stimulation (tDCS) is a non-invasive brain stimulation technique which was found to have a positive modulatory effect on online sequence acquisition or offline motor consolidation, depending on the relative role of the associated brain region. Primary motor regions (M1) and dorsolateral prefrontal cortices (DLPFC) have both been related to sequential learning. However, research so far did not systematically disentangle their differential roles in online and offline learning especially in more complex sequential paradigms. In this study, the influence of M1-tDCS and DLPFC-tDCS on complex sequential learning (online and offline) was investigated using a complex whole body serial reaction time task (CWB-SRTT) in 42 healthy volunteers. TDCS groups did not differ from sham tDCS group regarding their total time to complete the sequence and reaction time (online) and also not in terms of over-night consolidation (offline). Results may be related to unspecific parameters such as timing of the stimulation or current intensity but can also be attributed to the relative role of M1 and DLPFC during early complex learning. Future studies should consider investigating neural parameters during early complex CWB-SRTT learning to gain information on changes in neural activation within sequence acquisition with a specific focus on M1 and DLPFC.

## Introduction

Many activities in everyday living ranging from simple tasks such as getting ready for work to complex sports-related activities require performing several consecutive motor tasks in a sequential order. Most familiar motor sequences are performed in a highly automatized manner, but all have to be learned at the beginning. Usually, most familiar motor sequences are performed highly automatized, but injury, disease or even healthy aging can make active relearning necessary. To support successful learning, understanding mechanisms of sequential task acquisition/ consolidation and associated brain networks is crucial, especially since some motor sequences are established incidentally or implicit (Cleeremans, Destrebecqz, & Boyer, 1998). Sequential learning has been operationalized most frequently by using a serial reaction time task (SRTT (Nissen & Bullemer, 1987; Robertson, 2007)). SRTT learning is conceptualized by comparing responses to a repeating sequence of stimuli, which can be predicted and learned, with responses to randomly ordered stimuli, where only visuomotor associations but no temporal predictions are learned (Savic & Meier, 2016). Learning the SRTT involves motor, perceptual and declarative components (Robertson, 2007), thus a variety of different brain regions were found to be associated with it (Dayan & Cohen, 2011; Keele, Ivry, Mayr, Hazeltine, & Heuer, 2003), including frontal (Mizuguchi et al., 2019; Robertson, Tormos, Maeda, & Pascual-Leone, 2001), sensorimotor and cerebellar areas (Baldassarre, Filardi, Spadone, Penna, & Committeri, 2021; Doyon, Penhune, & Ungerleider, 2003; Orban et al., 2010) with their relative contribution depending on task complexity (Carey, Bhatt, & Nagpal, 2005; Gonzalez & Burke, 2018) and learning phase (Doyon & Benali, 2005).

Non-invasive brain stimulation techniques such as transcranial direct current stimulation (tDCS) have been widely used to investigate the causal relationship between stimulated brain regions and behavior (Hashemirad, Zoghi, Fitzgerald, & Jaberzadeh, 2016; Savic & Meier, 2016). In brief, tDCS modulates cortical excitability in a polarity-specific manner (Nitsche & Paulus, 2000) and, when applied over a longer period of time, also modulates cortical plasticity and behavior (Fritsch et al., 2010; Stagg, Bachtiar, & Johansen-Berg, 2011). TDCS can facilitate online sequence acquisition and offline motor consolidation, depending on the relative role of the associated brain region. During SRTT-learning, anodal tDCS (a-tDCS) over the primary motor cortex (M1) was mainly found to improve offline motor consolidation (Ehsani, Bakhtiary, Jaberzadeh, Talimkhani, & Hajihasani, 2016; Hashemirad et al., 2016; Krause, Meier, Dinkelbach, & Pollok, 2016; Reis et al., 2009), a process, usually noticeable as performance maintenance or improvement between sessions (Robertson, Pascual-Leone, & Miall, 2004). A positive effect of M1-stimulation on SRTT acquisition was only found in some studies (Kantak, Mummidisetty, & Stinear, 2012; Nitsche et al., 2003; Savic & Meier, 2016), while others did not find any a-tDCS effect during acquisition (Ehsani et al., 2016; Reis et al., 2009) or even reported worsening of performance (Keitel, Ofsteng, Krause, & Pollok, 2018). Besides, there is growing evidence for a substantial role of the prefrontal cortex (PFC) during sequential learning, at least in more complex learning paradigms (Galea, Albert, Ditye, & Miall, 2010; Meier et al., 2013; Mizuguchi et al., 2019). A-tDCS over the dorsolateral PFC (DLPFC) was found to improve implicit motor learning (Gladwin, den Uyl, & Wiers, 2012; Nakashima et al., 2021; Yamamoto, Ishii, Ishibashi, & Kohno, 2022). However, to our knowledge DLPFC-tDCS in SRTT learning has rarely been studied (Nakashima et al., 2021; Nitsche et al., 2003). Taken together, tDCS can facilitate SRTT sequence acquisition and consolidation, depending on the relative role of the associated brain region. M1 and DLPFC both substantially contribute to SRTT learning; however, research so far did not systematically disentangle their differential roles in online and offline SRTT learning especially in more complex paradigms.

Therefore, in the present study, we were aiming at investigating the relative contribution of DLPFC and M1 on sequential learning using a complex whole-body SRTT (CWB-SRTT). The CWB-SRTT was previously developed by our group (Maudrich, Kenville, Schempp, Noack, & Ragert, 2021; Mizuguchi et al., 2019) and is a modified version of a SRTT performed with the lower extremities. Therefore, it places higher demands on whole-body postural control and spatial orientation. Single-session tDCS over DLPFC or M1 was applied during CWB-SRTT acquisition and both online and offline gains were investigated using a two-day interventional paradigm. With regard to previous studies, we hypothesized, that (i) bilateral DLPFC-tDCS enhances online CWB-SRTT learning, indicated by faster completion time reduction in sequence trials compared to sham tDCS (Gladwin et al., 2012). Furthermore, we expected (ii) bilateral DLPFC-tDCS to enhance CWB-SRTT sequence specific learning (SSL), indicated by a time difference between sequence and random trials (Mizuguchi et al., 2019). For M1-tDCS, we mainly hypothesized (iii) a beneficial effect on motor consolidation, indicated by a between-session improvement or maintenance of completion time values.

## Material and Methods

### Ethical approval

This study was approved by the local ethics committee of Leipzig University (ref. nr. 191/19-ek). All participants provided written informed consent and all procedures were conducted in accordance with the Declaration of Helsinki.

### Participants

Recent studies investigating implicit motor learning using DLPFC-tDCS reported learning effects of medium size (Nakashima et al., 2021; Yamamoto et al., 2022; Zhu et al., 2015). Therefore, an a priori sample size calculation (G*Power 3.1) specified a total sample size of 15 participants to obtain a significant within-between interaction assuming a moderate effect size f = 0.25 and a power level of 1-β = 0.8 using a mixed model with three groups, fifteen measurement points, and a significance level of α < 0.05. For M1-tDCS, a robust medium effect for single session M1-tDCS on retention was reported (standardized mean difference: 0.71) in a meta-analysis investigating the effect of M1-tDCS on sequential learning (Hashemirad et al., 2016). Therefore, a priori sample size calculation (G*Power 3.1) specified a total sample size of 42 individuals to obtain a significant interaction assuming f = 0.25 and a power level of 0.8 using a mixed model with three groups and two measurement points (S15 TD1, S1 TD2). We enrolled a total of 42 volunteers in our experiment (14 per group, mean age: 24,29 ± 2,76 years, range: 20 – 31 years, 18 female). None of the participants had a history of neurological illness, and during the time of the experiment, no participant was taking any central-acting drugs. All participants were right-handed. Before and after the experimental procedure, all participants rated their levels of attention, fatigue and discomfort on a visual analog scale (VAS) to rule out unspecific effects of these factors on behavioral performance.

### Experimental procedure

Each participant performed two consecutive training sessions of a complex whole-body serial reaction time task (CWB-SRTT) with its lower extremity. On the first training day (TD1), tDCS was either applied (a) bilaterally over the left and right DLPFC (DLPFC-tDCS, group 1) or (b) over M1 leg area (M1-tDCS, group 2) or (c) as a sham intervention (s-tDCS, group 3), while participants performed a total number of 20 trials with four random trials (RT) at the beginning, a series of 15 sequence trials (ST) in the middle and one RT at the end of the session. Two RT were performed before tDCS was started to evaluate before-stimulation baseline performance, while two RT were performed during tDCS. Since CWB-SRTT performance took approximately 15 min, including 25-s inter-block intervals, tDCS administration was continued after task completion for approximately five minutes during rest at the end of the experimental session. On the second training day (TD2), participants performed 17 CWB-SRTT trials with one RT at the beginning,15 STs in between and one RT at the end without tDCS to evaluate potential effects on CWB-SRTT skill consolidation and skill learning development.

### Sensorimotor task: whole-body serial reaction time task (CWB-SRTT)

A detailed description of the four-directional CWB-SRTT can be found in previous publications of our group (Maudrich et al., 2021; Mizuguchi et al., 2019). In brief, participants stood on two center plates, which were surrounded by a four-directional plate array. Participants were instructed to look at a monitor in front of them, which displayed the target cue in one of four squares, corresponding to the four surrounding plates and respond to the cue by stepping onto the corresponding plate as quickly as possible after its appearance. Left foot usage was instructed for left-side plates and right foot usage for right-side plates. After each response, participants were asked to move back to their initial position, where 500ms later the next cue appeared. Target cues remained visible until the correct response was made and incorrect responses were indicated by a red flashing on the monitor. In total, 12 items per trial were presented. ST included the following cue order: 2-3-2-4-1-3-1-4-3-4-2-1 (1: front left, 2: front right, 3: back left, 4: back right), thus providing a balanced number of movements to each of the four directions. All participants were naïve regarding the learning sequence and were explicitly not instructed about the presence of a sequence to maximize the possibility of performing an implicit motor learning task. To evaluate explicit sequence awareness, sequence knowledge was tested after the second training day by asking the participant (a) whether they recognized a sequential pattern in the visual cue order and (b) to recall as many elements of the sequence as possible without external help. RT included a pseudo random cue order with equal probabilities regarding each number and a limit of maximally three consecutive repetitions per item. Each block lasted for a maximum of 25-s and between blocks a 25-s interblock-interval was included. After each trial, participants received feedback about the total time they took to complete the sequence. A custom-made script operated the CWB-SRTT (C#, Microsoft Visual Studio 2017).

### Transcranial direct current stimulation

Transcranial direct current stimulation (tDCS) was applied via a battery-driven stimulator (neuroConn GmbH, Ilmenau, Germany) with two attached electrodes. In case of DLPFC stimulation (DLPFC-tDCS), anodal and cathodal electrodes both had a size of 5×7 cm and were positioned to produce a bilateral stimulation pattern (DaSilva et al., 2015). Thus, according to the 10-20 system, the cathode was positioned over F3 (MNI-Coordinate: x = -34, y = 26, z = 44 (Keeser et al., 2011), while the anode was positioned over F4 (MNI-Coordinate: x = 34, y = 26, z = 44). TDCS of 2 mA was applied since previous research found PFC modulation using 2 mA tDCS, but not smaller (Boggio et al., 2006; Teo, Hoy, Daskalakis, & Fitzgerald, 2011). For M1 leg area stimulation (M1-tDCS), a 5×7 cm anode was placed over the leg area of M1 (coordinates: x = 0, y = -24, z = 75, (Long, Goltz, Margulies, Nierhaus, & Villringer, 2014; Taubert, Mehnert, Pleger, & Villringer, 2016), while a 5×10 cm cathode was placed over the supraorbital area. Anatomical landmarks were identified using neuronavigation (Localite TMS-Navigator, Bonn, Germany) using our predefined MNI coordinates. After localization with the neuronavigation system, the scalp of the participants was first rubbed with alcohol, then both electrodes were soaked in saline and fixed to the scalp with rubber bands. tDCS was applied with a current intensity of 2 mA. Current was applied for 20 min during CWB-SRTT task performance with a fade-in and fade-out period of each 30 s each. During sham stimulation (s-tDCS), the current was ramped up for 30 s, held constant at 1 mA for 30 s, and ramped down for 30 s. This short duration of stimulation has been shown to elicit no changes in cortical excitability while it may provide the same tingling sensation on the scalp of the participant (Nitsche et al., 2008). Generally, the impedance was monitored and kept under 10 kΩ (average: 3.87 ± 4.96 10 kΩ). Both researcher and participants were blinded.

### tDCS current flow simulation

Electric field distributions were simulated based on a finite element model of a representative head using the open-source SimNIBS software (Thielscher, Antunes, & Saturnino, 2015) to approximate current flow. For (a) bilateral DLPFC-tDCS, anode and cathode both were defined according to our anatomical landmarks (anode: 34, y = 26, z = 44, cathode: x = -34, y = 26, z = 44), each with a size of 7 × 5 cm. For (b) M1-tDCS, anode was defined according using predefined coordinates (x = 0, y = -24, z = 75), while a 5×10 cm cathodes center location was established at Fpz. For both stimulation conditions (a) and (b), a current intensity of 2 mA was selected. For bilateral DLPFC-tDCS, maximum electrical field strength (0.422 V/m) was determined between the two electrodes, covering both dorsolateral and medial prefrontal cortices. For M1-tDCS, maximum field strength (0.354 V/m) was found anterior to the anode, corresponding to bilateral leg motor cortices but also premotor and dorsolateral prefrontal cortices (see Fig. 2 for details).

**Figure 1:**
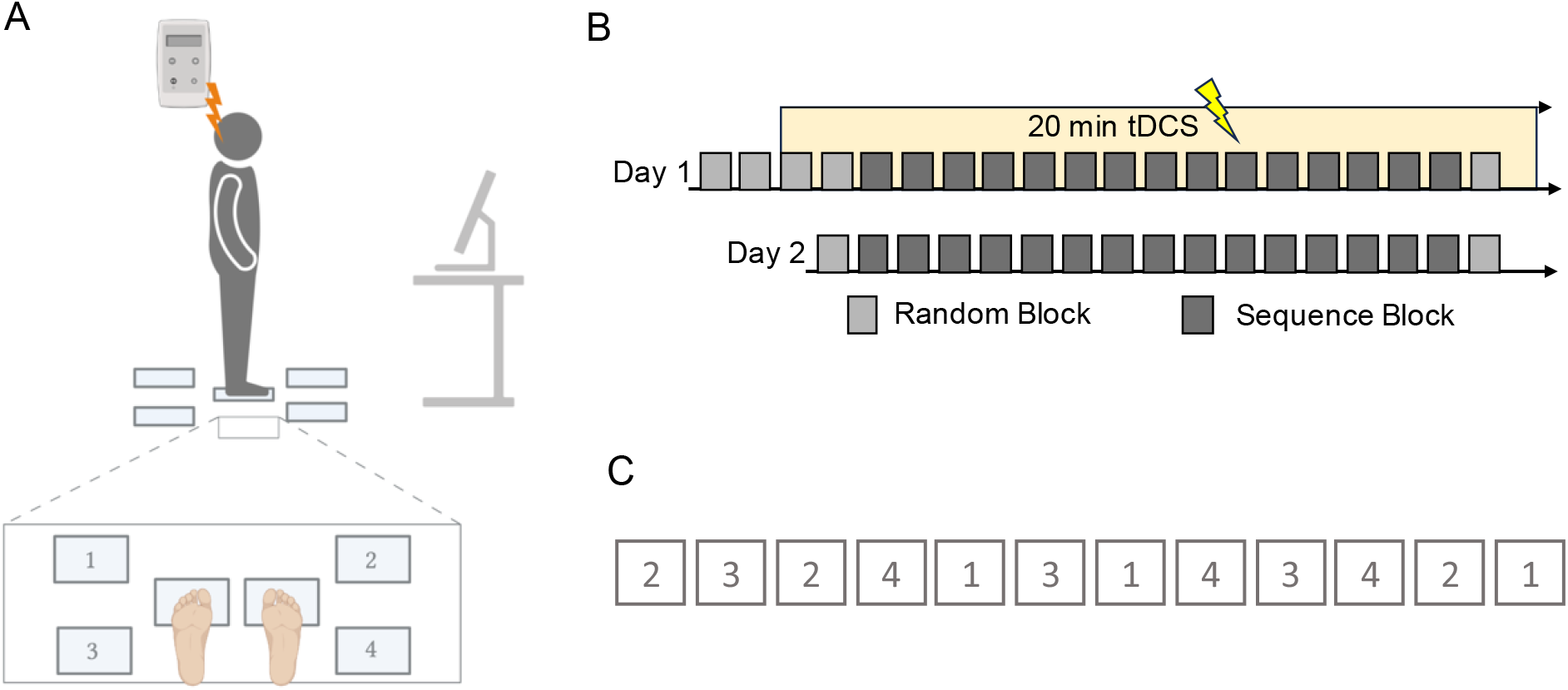
Complex Whole-body serial reaction time task (CWB-SRTT). **A:** experimental setup: participants received 20 minutes of tDCS, while they performed the CWB-SRTT. Created with BioRender.com (2023). **B:** time line experimental setup. **C:** Sequential elements (N=12). Number corresponds to plate number

**Figure 2:**
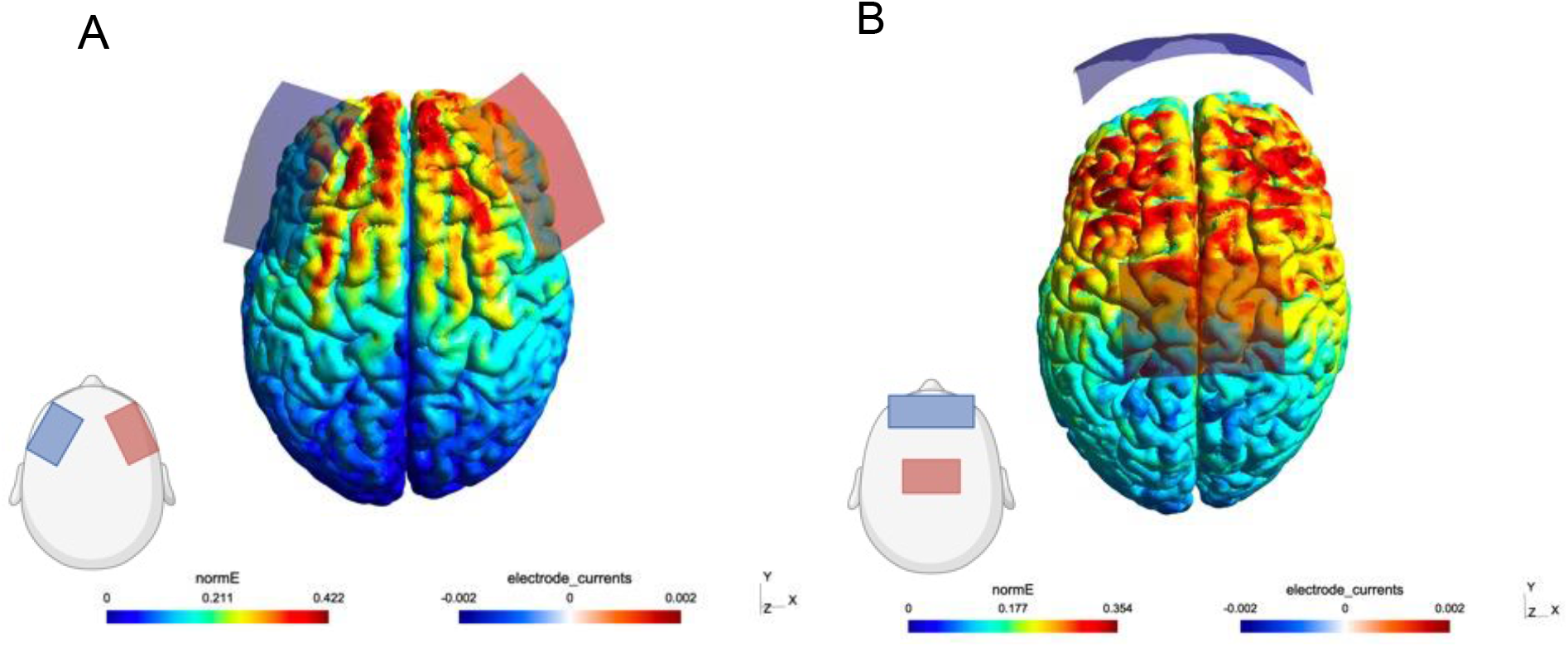
Transcranial direct current stimulation (tDCS) current flow simulation. Normalized electrical field strength (V/m) is depicted for **A:** bilateral DLPFC-tDCS and **B:** primary motor cortex (M1)-tDCS covering the leg motor cortex. Normalized electrical field strength (V/m) is indicated through colormaps with blue representing lowest and red representing highest field strengths, respectively. Current flow image was created using the SIMNIBS software version 3.2. (Thielscher et al., 2015).

**Figure 3:**
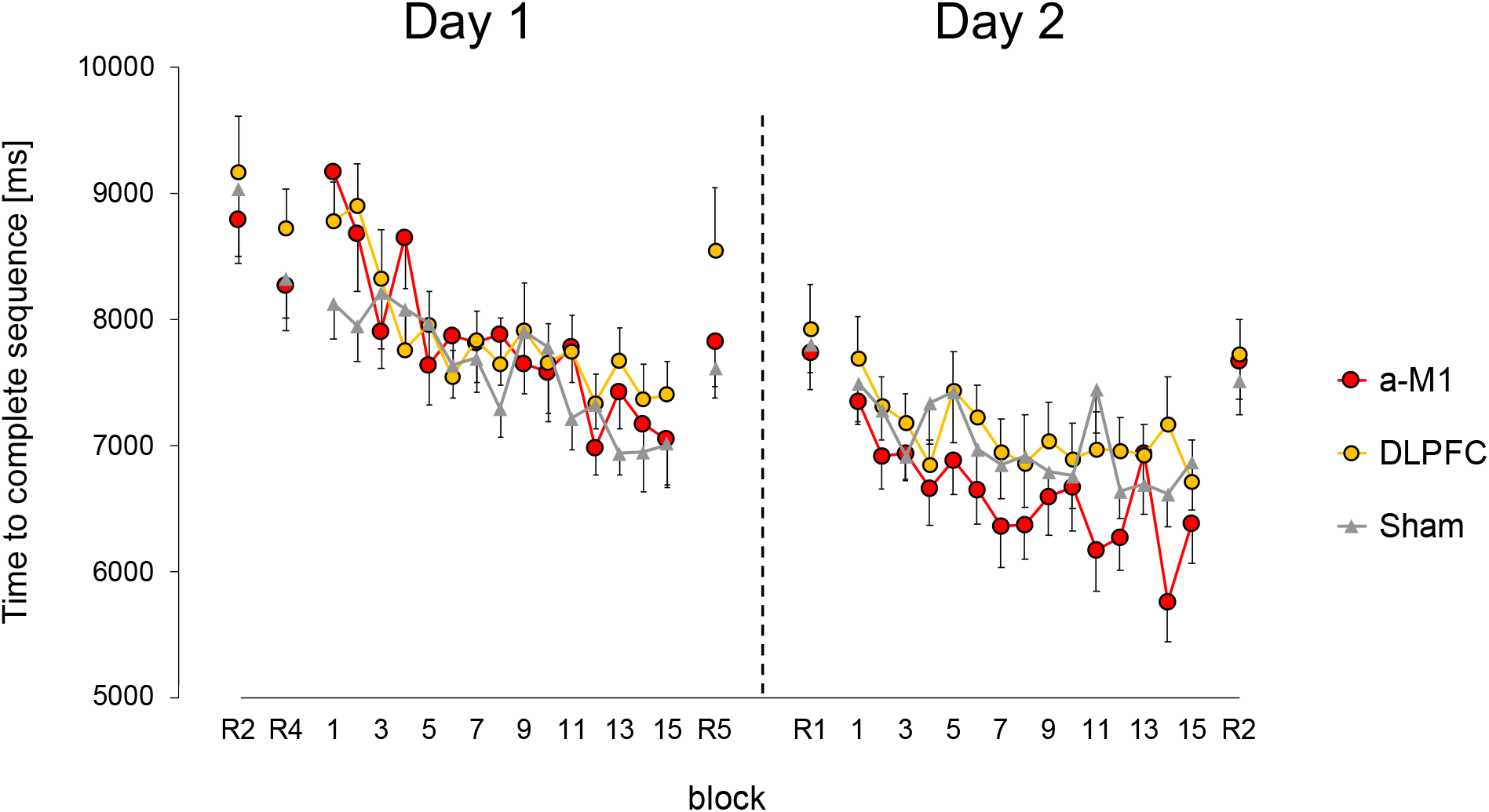
Time to complete sequence (TCS). Line Graph shows average values ± standard error. Red line corresponds to a-M1-tDCS, yellow line to DLPFC-tDCS, grey line represents sham-tDCS group. Dotted line represents separation of training days. DLPFC: bilateral dorsolateral prefrontal cortex stimulation, a-M1: M1 leg area stimulation.

**Figure 4:**
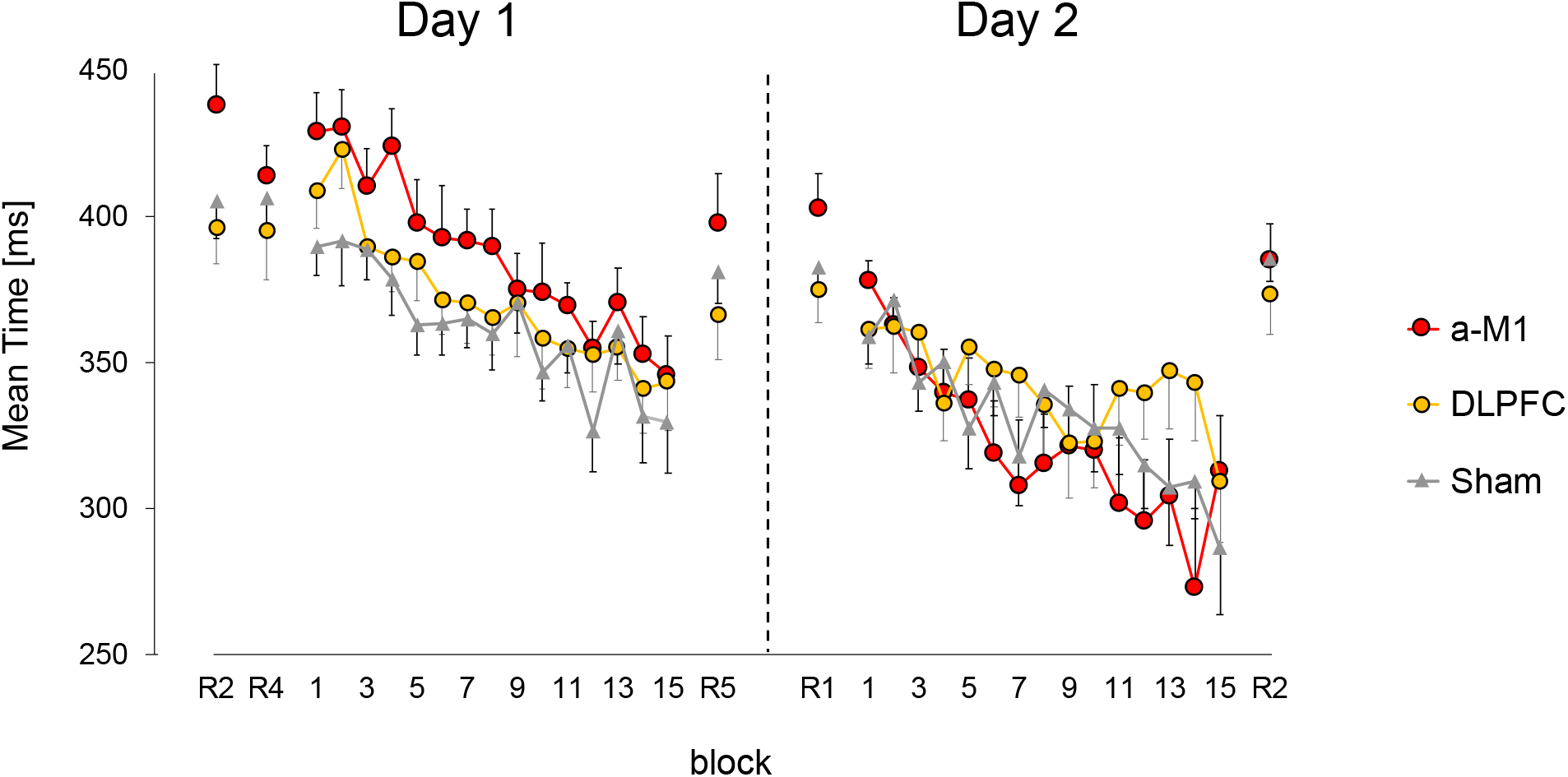
Mean Reaction Time (ReT). Line graphs display average values and standard deviation. A-M1-tDCS is represented by the red line, DLPFC-tDCS by the yellow line, and the sham-tDCS group is shown by the grey line. Training days are separated by a dotted line. DLPFC: bilateral dorsolateral prefrontal cortex stimulation, a-M1: M1 leg area stimulation.

### Data Analyses

All CWB-SRTT trials with a duration >1500ms were removed from the analysis since we do not expect them to represent physiological responses. Furthermore, all trials with errors were removed. CWB-SRTT learning was classified as the total time participants took to complete the sequence (*time to complete sequence*: TCS). Furthermore, our CWB-SRTT setting allowed TCS differentiation into reaction and movement time. Reaction time (ReT) was operationalized by classifying the time difference between onset of the visual stimulus and response foot lifting from one of the middle plates. Movement time (MoT) was operationalized by examining the time difference between response foot lifting from the middle plate and contact on one of the target plates. Signals were recorded at a sampling rate of 1000 Hz. We calculated the median of ReT and MoT, blockwise for each participant. Since ReT was the main outcome parameter, previous studies found to be associated with CWB-SRTT learning (Maudrich et al., 2021; Mizuguchi et al., 2019), we averaged ReT for each participant, resulting in 18 relevant ReT on TD1 per individual with second (R2), fourth (R4) and last sequence ReT (R5) considered as RT and ReT five to nineteen (S1 - S15) considered as ST. R1 and R3 were not included in the analysis since they were considered practice or habituation trials. On TD 2, only one RT was captured before sequential learning (R1), followed by 15 ST (S1-S15) and one RT (R2) in the end. Sequence specific learning (SSL) was evaluated by calculating Δ_ReT_ between R5 and S15. General non-sequence-specific learning (GL) was evaluated by calculating Δ_ReT_ between R5 and R2. Explicit sequence knowledge was evaluated by (a) counting the number of “yes” answers per group and (b) by summing the number of correctly recalled consecutive items of the learning sequence.

### Statistical Analyses

Statistical analyses were performed using JASP (Version 0.16, JASP Team 2021). The normality of the data was assessed by Shapiro-Wilk testing (α = 0.05). To compare initial performance between groups, TCS and ReT at R2 were evaluated using one-way ANOVAs. Skill Learning was assessed within and between groups by evaluating TCS and ReT differences on TD1 and TD2 using RM-ANOVAs with between-subject factor GROUP (DLPFC-tDCS, M1-tDCS, s-tDCS) and within-subject factor SEQUENCE (S1-S15). Consolidation was evaluated by comparing ReT performance on S15 TD1 and S2 TD2 using an RM-ANOVA with factor GROUP and TIME. Greenhouse-Geisser correction was implemented in case of sphericity violation. SSL and GL were evaluated separately for TD1 and TD2 using one-way ANOVAs. Equal properties between groups regarding sequence awareness were tested using a multinomial χ^2^ Test. Number of recalled items were compared between groups using a one-way ANOVA. The statistical threshold for all analyses was set at p < 0.05. Effect sizes were expressed using partial eta squared (η_p_^2^) for ANOVAs.

## Results

### Demographics

Demographic variables did not differ between groups (see Table 1 for details). All participants tolerated the stimulation well and none reported any unexpected side effects from tDCS stimulation. A multinominal test revealed no difference between members of the DLPFC-tDCS, M1-tDCS and s-tDCS group in the ability to judge their group belonging (χ ^2^ (3) = 1.16, p = .76), indicating that blinding of conditions was effective.

**Table 1:** TDCS groups did not differ regarding age (ANOVA, F(2,39)=1.102, p=.34), number of female participants (multinominal test: χ^2^(3) = 1.17, p = .76) and hours of sports per week (ANOVA, F(2,39)=.36, p=.7). Furthermore, changes in attention (A), fatigue (F) and discomfort (D) levels (**Δ** A, F, P TD1) on TD1 did not differ between groups (RM-ANOVAs, interaction effect, A: F(2,39)=.56, p=.58, F: F(2,39)=1.72, p=.19, P: F(2,39)=.49, p=.62).

| Group | Age (years) | Gender (f/m) | H Sports/week | $\Delta$ A TD1 | $\Delta$ F TD1 | $\Delta$ D TD1 |
| --- | --- | --- | --- | --- | --- | --- |
| DLPFC-tDCS | 23.43 ± 2.6 | 7/7 | 7.25 ± 2.73 | -0.18 ± 0.77 | -0.11 ± 1.11 | -0.14 ± 0.36 |
| M1-tDCS | 24.93 ± 3.3 | 6/8 | 8.04 ± 4.78 | -0.53 ± 1.37 | -0.79 ± 1.25 | 0.07 ± 0.27 |
| S-tDCS | 24.5 ± 2.1 | 5/9 | 6.71 ± 3.61 | -0.57 ± 1.04 | -0.21 ± 0.7 | -0.07 ± 0.27 |

### Time to complete sequence

Initial baseline TCS performance on R2 before tDCS onset showed no significant difference between groups (F(39) = .18, p = .83, η_p_^2^ = .009), indicating no group difference prior to the stimulation. We found a significant main effect of time for TCS across groups both on TD1 (F(9.46, 369.01) = 10.48, p < .001, η_p_^2^ = .13) and on TD 2 (F(8.96, 349.48) = 4.86, p <. 001, η_p_^2^ = .11), indicating that all participants substantially reduced their TCS over time. We found no significant difference in TCS between groups across all trials on TD1 (F(2,39) = .41, P = .67, η_p_^2^ = .02) and TD2 (F(2,39) = 1.52, p = .232, η_p_^2^ = .07) and also no significant group difference in TCS reduction over time on TD1 (F(18.92, 369.01) = 1.23, p = .23, η_p_^2^ = .06) or TD2 (F(17.92, 349.38) = 1.28, p = .2, η_p_^2^ = .06).

### Reaction Time (ReT)

ReT performance on R2 did not differ between groups (F(2,39) = 2.83, p = .07, η_p_^2^ = .13). Analog to TCS, we also found ReT to be substantially reduced over time across groups on TD 1 (F(7.79, 303.73) = 19.63, p < .001, η_p_^2^ = .34) and TD 2 (F(6.36, 247.98) = 9.11, p< .001, η_p_^2^ = .19). Groups did not differ in ReT performance (TD1: F(2,39) = 1.78, p = .18, η_p_^2^ = .08, TD2: F(2,39) = 0.58, p = .57, η_p_^2^ = .03) and ReT reduction over time (TD1: F(15.58, 303.73) = 0.67, p = .82, η_p_^2^ = .03, TD2: F(12.72, 247.98) = 1.75, p = .054, η_p_^2^ = .08). Furthermore, consolidation was similar in all groups (F(2,39) = .24, p = .79, η_p_^2^ = .01), ReT increased significantly (F(2,39) = 8.81, p = .005, η_p_^2^ = .18), on average by 26.16 ± 55.11 ms, from S15 TD1 to S1 TD2.

### Sequence Specific Learning (SSL) and General Learning (GL)

When comparing SSL between groups, we found no significant difference on TD 1 (F(2,39) = 1.03, p = .37, η_p_^2^ = .05, see also Figure 5), TD 2 (F(2,39) = .76, p = .47, η_p_^2^ = .04) and also total SSL learning did not differ between groups (F(2,39) = 1.09, p = .35, η_p_^2^ = .05). GL did not differ between groups, either (TD1: F(2,39) = .41, p = .67, η_p_^2^ = .02, TD2: F(2,39) = .92, p = .41, η_p_^2^ = .05, total learning (F(2,39) = 1.39, p = .26, η_p_^2^ = .07).

**Figure 5:**
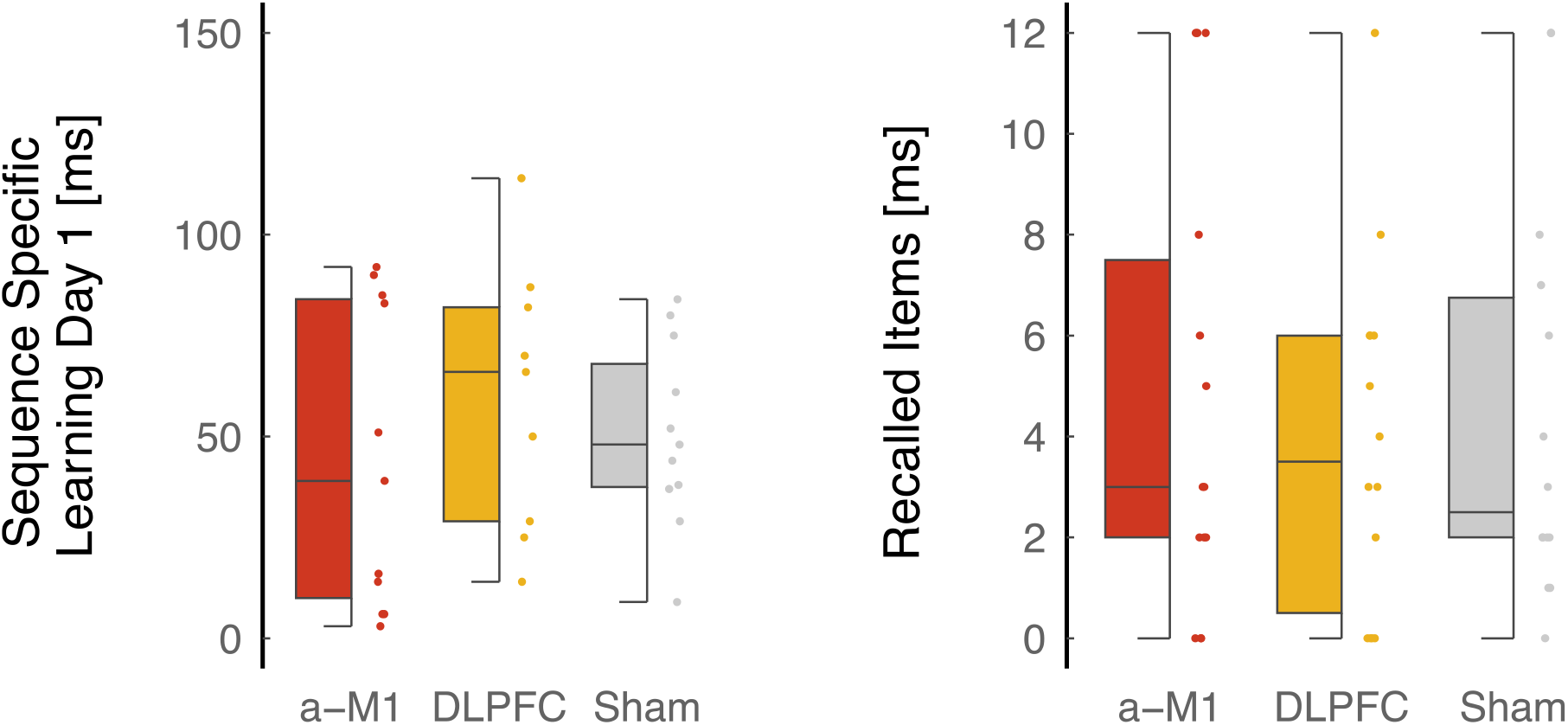
Sequence Specific Learning (SSL) TD 1 and Number of recalled items (NoI). Boxplots show medians with 25th and 75th percentiles, Dot plots showing individual values. Outliers were excluded. Red box corresponds to M1-tDCS, yellow boxes represent DLPFC-tDCS, grey boxes represent sham tDCS.

### Explicit Sequence Knowledge

Equal properties between groups regarding sequence awareness were found (multinomial χ^2^(3) = 1.16, p = .76). Furthermore, the number of items (NoI), participants were able to recall after the end of TD2, did not differ between groups (F(2,39) = .16, p = .85, η_p_^2^ = .01, see also Figure 5).

## Discussion

The main aim of the current study was to investigate the relative contribution of DLPFC and M1 on sequential learning using a complex whole-body SRTT (CWB-SRTT). Contrary to our hypotheses, DLPFC-tDCS did not enhance online CWB-SRTT learning. More precisely, we couldn’t confirm hypothesis (i) and (ii) but found similar TCS and ReT reduction in all groups and also comparable SSL ratios. These findings contradict previous studies which suggested improved sequence acquisition associated with DLPFC-tDCS (Nakashima et al., 2021; Yamamoto et al., 2022; Zhu et al., 2015). Nakashima et al. (2021) found reaction time reductions associated with DLPFC stimulation, however, effects were only found post stimulation. In the current study, post stimulation effects were not investigated. Instead, tDCS outlasted the SRTT training period in all cases, thus, one can only speculate about reaction time decreases after termination of the stimulation. Furthermore, in previous studies, DLPFC-tDCS modulated sequential learning only in motor tasks where the supervisory attention system (SAS) was involved but not in motor tasks without SAS guidance (Yamamoto et al., 2022). In the current study, no explicit experimental manipulation of attentional parameters was done, however our VAS scale assessment revealed no substantial increase of attention throughout the experimental session. It seems reasonable to assume that CWB-SRTT learning does not require SAS guidance, which is why potential DLPFC-tDCS-induced enhancement of attentional parameters may not have resulted in superior CWB-SRTT motor learning performance. Moreover, Zhu et al. (2015) found implicit motor learning enhancement after cathodal tDCS was applied over the left DLPFC, which was argued by a shift in dominance from explicit to implicit memory system via reduction of explicit verbal-analytical involvement in movement control (Zhu et al., 2015). Our current flow simulation showed that bilateral DLPFC-tDCS resulted in a widespread activation increase covering large amounts of both frontal cortices. High field strength in both left and right DLPFC may have resulted in more explicit memory system usage. However, our parameter of explicit task knowledge (NoI) did not differ between groups, indicating that at least explicit CWB-SRTT learning was not specifically supported by bilateral DLPFC-tDCS. While enhancement of implicit motor learning was mainly associated with left DLPFC stimulation (Nakashima et al., 2021; Yamamoto et al., 2022; Zhu et al., 2015), little is known about right DLPFC activation and its role in motor learning subsystems. If bilateral DLPFC-tDCS activated both explicit and implicit memory systems, simultaneously, it may have resulted in a conflicting situation of both memory systems (Cohen & Robertson, 2011), thus inducing no benefit in motor learning.

We also could not confirm hypothesis (iii), assuming a beneficial effect of M1-tDCS on motor consolidation. M1-tDCS effects on motor consolidation have been shown in a variety of different motor settings but usually using the upper limb (Hashemirad et al., 2016). However, stimulating M1 of the lower limb is more complex given the anatomical location of the leg motor cortex (Madhavan, Sriraman, & Freels, 2016; Rezaee & Dutta, 2018). Therefore, transferring upper-limb findings to lower-limb findings may not meet all challenges associated with lower limb M1-tDCS and lower limb M1-tDCS may not be as effective as upper-limb tDCS in area targeting and producing associated behavioral improvements. Nonetheless, there are also studies confirming beneficial effects of lower limb M1-tDCS even in complex motor settings (Devanathan & Madhavan, 2016; Kaminski et al., 2016). Other studies found no effect on reaction time using lower limb M1-tDCS (Seidel & Ragert, 2019). Thus, not in all cases, M1-tDCS was shown to be beneficial for learning and its relative effect may depend on the nature of the motor task. As already mentioned, CWB-SRTT can be considered a complex motor learning task with additional requirements to the postural system since the task is performed during standing. Furthermore, rear plates were invisible to the participant during task performance, thus, a relatively precise internal model about the relative plate distance and associated stride length was needed to successfully meet the plates. Internal model creation and organization was mainly found to be acquired in the cerebellum (Imamizu, Kuroda, Miyauchi, Yoshioka, & Kawato, 2003; Imamizu et al., 2000), therefore, CWB-SRTT learning could be more related to cerebellar activation. Future studies could evaluate how cerebellar modulation interacts with CWB-SRTT learning. Additionally, even though previous studies did find a beneficial effect of M1-tDCS on SRTT learning, only effects on TCS were investigated (Savic & Meier, 2016). CWB-SRTT analyses additionally allows disentangling reaction and movement time, thus making a precise interpretation of cognitive and motor mechanisms of learning possible. CWB-SRTT learning was mainly found to be associated with ReT reductions while MoT did not systematically decrease over time (Mizuguchi et al., 2019). Therefore, one can conclude that mainly cognitive processes are responsible for CWB-SRTT learning, which is one potential explanation that M1-tDCS did not result in superior CWB-SRTT performance. Furthermore, another study used 4 mA current intensity to robustly modify sequential learning using M1-tDCS (Hsu, Shereen, Cohen, & Parra, 2023), thus one could speculate that our current intensity was not sufficient to target M1 and induce a robust behavioral improvement.

Our study faces some limitations. One major issue of tDCS is its relatively low spatial focality (Buch et al., 2017), resulting in widespread co-activation of adjacent but also functionally connected brain regions. Our current flow model shows that during both DLPFC and M1 stimulation, large parts of the frontal cortex including pre-motor cortex, supplementary motor area and prefrontal regions were stimulated, even though the strongest electrical field was found anterior to the active electrode. Previous studies with similar electrode positions found tDCS-induced behavioral improvements (DaSilva et al., 2015; Kaminski et al., 2016), thus one cannot conclude that widespread cortical activation is not efficient. However, during CWB-SRTT learning, activation of brain regions with potentially complementary functions may have resulted in a conflicting situation, not allowing relevant areas to determine the learning process. It can be stressed that isolated activation of a single hemisphere region was not measured in the current study and therefore, potentially also behavioral effects are missing. Another limitation of the current study was that no additional brain imaging was performed. Joint activation increases or decreases of brain regions induced by tDCS and movement-induced neural activation or deactivation may have resulted in a specific brain activation pattern not displayable given our current study design. We therefore strongly recommend future studies to additionally determine brain activation changes, best during task execution. The fact that the direction of stimulation (excitatory vs. inhibitory) depends on the direction of current flow in the brain, which may differ across persons and between brain areas due to anatomical differences, is another drawback of tDCS (Buch et al., 2017).

Taken together, our study shows that tDCS application during complex sequential motor task acquisition is possible. However, our main aim was to disentangle roles of DLPFC and M1 during early complex sequential learning. We could not confirm a performance-enhancing effect of tDCS neither on online nor offline learning parameters. Results may be related to unspecific parameters such as timing of the stimulation or current intensity but can also be attributed to the relative role of M1 and DLPFC during early complex learning. Future studies should consider investigating neural parameters during early complex CWB-SRTT learning to gain information on changes in neural activation within sequence acquisition with a specific focus on M1 and DLPFC.

## References

Baldassarre, A., Filardi, M. S., Spadone, S., Penna, S. D., & Committeri, G. (2021). Distinct connectivity profiles predict different in-time processes of motor skill learning. Neuroimage, 238, 118239. doi:10.1016/j.neuroimage.2021.118239

Boggio, P. S., Castro, L. O., Savagim, E. A., Braite, R., Cruz, V. C., Rocha, R. R., … Fregni, F. (2006). Enhancement of non-dominant hand motor function by anodal transcranial direct current stimulation. Neurosci Lett, 404(1-2), 232–236. doi:10.1016/j.neulet.2006.05.051

Buch, E. R., Santarnecchi, E., Antal, A., Born, J., Celnik, P. A., Classen, J., … Cohen, L. G. (2017). Effects of tDCS on motor learning and memory formation: A consensus and critical position paper. Clin Neurophysiol, 128(4), 589–603. doi:10.1016/j.clinph.2017.01.004

Carey, J. R., Bhatt, E., & Nagpal, A. (2005). Neuroplasticity promoted by task complexity. Exerc Sport Sci Rev, 33(1), 24–31. Retrieved from https://www.ncbi.nlm.nih.gov/pubmed/15640717

Cleeremans, A., Destrebecqz, A., & Boyer, M. (1998). Implicit learning: news from the front. Trends Cogn Sci, 2(10), 406–416. doi:10.1016/s1364-6613(98)01232-7

Cohen, D. A., & Robertson, E. M. (2011). Preventing interference between different memory tasks. Nat Neurosci, 14(8), 953–955. doi:10.1038/nn.2840

DaSilva, A. F., Truong, D. Q., DosSantos, M. F., Toback, R. L., Datta, A., & Bikson, M. (2015). State-of-art neuroanatomical target analysis of high-definition and conventional tDCS montages used for migraine and pain control. Front Neuroanat, 9, 89. doi:10.3389/fnana.2015.00089

Dayan, E., & Cohen, L. G. (2011). Neuroplasticity subserving motor skill learning. Neuron, 72(3), 443–454. doi:10.1016/j.neuron.2011.10.008

Devanathan, D., & Madhavan, S. (2016). Effects of anodal tDCS of the lower limb M1 on ankle reaction time in young adults. Exp Brain Res, 234(2), 377–385. doi:10.1007/s00221-015-4470-y

Doyon, J., & Benali, H. (2005). Reorganization and plasticity in the adult brain during learning of motor skills. Curr Opin Neurobiol, 15(2), 161–167. doi:10.1016/j.conb.2005.03.004

Doyon, J., Penhune, V., & Ungerleider, L. G. (2003). Distinct contribution of the cortico-striatal and cortico-cerebellar systems to motor skill learning. Neuropsychologia, 41(3), 252–262. doi:10.1016/s0028-3932(02)00158-6

Ehsani, F., Bakhtiary, A. H., Jaberzadeh, S., Talimkhani, A., & Hajihasani, A. (2016). Differential effects of primary motor cortex and cerebellar transcranial direct current stimulation on motor learning in healthy individuals: A randomized double-blind sham-controlled study. Neurosci Res, 112, 10–19. doi:10.1016/j.neures.2016.06.003

Fritsch, B., Reis, J., Martinowich, K., Schambra, H. M., Ji, Y., Cohen, L. G., & Lu, B. (2010). Direct current stimulation promotes BDNF-dependent synaptic plasticity: potential implications for motor learning. Neuron, 66(2), 198–204. doi:10.1016/j.neuron.2010.03.035

Galea, J. M., Albert, N. B., Ditye, T., & Miall, R. C. (2010). Disruption of the dorsolateral prefrontal cortex facilitates the consolidation of procedural skills. J Cogn Neurosci, 22(6), 1158–1164. doi:10.1162/jocn.2009.21259

Gladwin, T. E., den Uyl, T. E., & Wiers, R. W. (2012). Anodal tDCS of dorsolateral prefontal cortex during an Implicit Association Test. Neurosci Lett, 517(2), 82–86. doi:10.1016/j.neulet.2012.04.025

Gonzalez, C. C., & Burke, M. R. (2018). Motor Sequence Learning in the Brain: The Long and Short of It. Neuroscience, 389, 85–98. doi:10.1016/j.neuroscience.2018.01.061

Hashemirad, F., Zoghi, M., Fitzgerald, P. B., & Jaberzadeh, S. (2016). The effect of anodal transcranial direct current stimulation on motor sequence learning in healthy individuals: A systematic review and meta-analysis. Brain Cogn, 102, 1–12. doi:10.1016/j.bandc.2015.11.005

Hsu, G., Shereen, A. D., Cohen, L. G., & Parra, L. C. (2023). Robust enhancement of motor sequence learning with 4 mA transcranial electric stimulation. Brain Stimul, 16(1), 56–67. doi:10.1016/j.brs.2022.12.011

Imamizu, H., Kuroda, T., Miyauchi, S., Yoshioka, T., & Kawato, M. (2003). Modular organization of internal models of tools in the human cerebellum. Proc Natl Acad Sci U S A, 100(9), 5461–5466. doi:10.1073/pnas.0835746100

Imamizu, H., Miyauchi, S., Tamada, T., Sasaki, Y., Takino, R., Putz, B., … Kawato, M. (2000). Human cerebellar activity reflecting an acquired internal model of a new tool. Nature, 403(6766), 192–195. doi:10.1038/35003194

Kaminski, E., Steele, C. J., Hoff, M., Gundlach, C., Rjosk, V., Sehm, B., … Ragert, P. (2016). Transcranial direct current stimulation (tDCS) over primary motor cortex leg area promotes dynamic balance task performance. Clin Neurophysiol, 127(6), 2455–2462. doi:10.1016/j.clinph.2016.03.018

Kantak, S. S., Mummidisetty, C. K., & Stinear, J. W. (2012). Primary motor and premotor cortex in implicit sequence learning--evidence for competition between implicit and explicit human motor memory systems. Eur J Neurosci, 36(5), 2710–2715. doi:10.1111/j.1460-9568.2012.08175.x

Keele, S. W., Ivry, R., Mayr, U., Hazeltine, E., & Heuer, H. (2003). The cognitive and neural architecture of sequence representation. Psychol Rev, 110(2), 316–339. doi:10.1037/0033-295x.110.2.316

Keeser, D., Meindl, T., Bor, J., Palm, U., Pogarell, O., Mulert, C., … Padberg, F. (2011). Prefrontal transcranial direct current stimulation changes connectivity of resting-state networks during fMRI. J Neurosci, 31(43), 15284–15293. doi:10.1523/JNEUROSCI.0542-11.2011

Keitel, A., Ofsteng, H., Krause, V., & Pollok, B. (2018). Anodal Transcranial Direct Current Stimulation (tDCS) Over the Right Primary Motor Cortex (M1) Impairs Implicit Motor Sequence Learning of the Ipsilateral Hand. Front Hum Neurosci, 12, 289. doi:10.3389/fnhum.2018.00289

Krause, V., Meier, A., Dinkelbach, L., & Pollok, B. (2016). Beta Band Transcranial Alternating (tACS) and Direct Current Stimulation (tDCS) Applied After Initial Learning Facilitate Retrieval of a Motor Sequence. Front Behav Neurosci, 10, 4. doi:10.3389/fnbeh.2016.00004

Long, X., Goltz, D., Margulies, D. S., Nierhaus, T., & Villringer, A. (2014). Functional connectivity-based parcellation of the human sensorimotor cortex. Eur J Neurosci, 39(8), 1332–1342. doi:10.1111/ejn.12473

Madhavan, S., Sriraman, A., & Freels, S. (2016). Reliability and Variability of tDCS Induced Changes in the Lower Limb Motor Cortex. Brain Sci, 6(3). doi:10.3390/brainsci6030026

Maudrich, T., Kenville, R., Schempp, C., Noack, E., & Ragert, P. (2021). Comparison of whole-body sensorimotor skill learning between strength athletes, endurance athletes and healthy sedentary adults. Heliyon, 7(8), e07723. doi:10.1016/j.heliyon.2021.e07723

Meier, B., Weiermann, B., Gutbrod, K., Stephan, M. A., Cock, J., Muri, R. M., & Kaelin-Lang, A. (2013). Implicit task sequence learning in patients with Parkinson’s disease, frontal lesions and amnesia: The critical role of fronto-striatal loops. Neuropsychologia, 51(14), 3014–3024. doi:10.1016/j.neuropsychologia.2013.10.009

Mizuguchi, N., Maudrich, T., Kenville, R., Carius, D., Maudrich, D., Villringer, A., & Ragert, P. (2019). Structural connectivity prior to whole-body sensorimotor skill learning associates with changes in resting state functional connectivity. Neuroimage, 197, 191–199. doi:10.1016/j.neuroimage.2019.04.062

Nakashima, S., Koeda, M., Ikeda, Y., Hama, T., Funayama, T., Akiyama, T., … Okubo, Y. (2021). Effects of anodal transcranial direct current stimulation on implicit motor learning and language-related brain function: An fMRI study. Psychiatry Clin Neurosci, 75(6), 200–207. doi:10.1111/pcn.13208

Nissen, M. J., & Bullemer, P. (1987). Attentional Requirements of Learning - Evidence from Performance-Measures. Cognitive Psychology, 19(1), 1–32. doi:Doi 10.1016/0010-0285(87)90002-8

Nitsche, M. A., Cohen, L. G., Wassermann, E. M., Priori, A., Lang, N., Antal, A., … Pascual-Leone, A. (2008). Transcranial direct current stimulation: State of the art 2008. Brain Stimul, 1(3), 206–223. doi:10.1016/j.brs.2008.06.004

Nitsche, M. A., & Paulus, W. (2000). Excitability changes induced in the human motor cortex by weak transcranial direct current stimulation. J Physiol, 527 Pt 3, 633–639. doi:10.1111/j.1469-7793.2000.t01-1-00633.x

Nitsche, M. A., Schauenburg, A., Lang, N., Liebetanz, D., Exner, C., Paulus, W., & Tergau, F. (2003). Facilitation of implicit motor learning by weak transcranial direct current stimulation of the primary motor cortex in the human. J Cogn Neurosci, 15(4), 619–626. doi:10.1162/089892903321662994

Orban, P., Peigneux, P., Lungu, O., Albouy, G., Breton, E., Laberenne, F., … Doyon, J. (2010). The multifaceted nature of the relationship between performance and brain activity in motor sequence learning. Neuroimage, 49(1), 694–702. doi:10.1016/j.neuroimage.2009.08.055

Reis, J., Schambra, H. M., Cohen, L. G., Buch, E. R., Fritsch, B., Zarahn, E., … Krakauer, J. W. (2009). Noninvasive cortical stimulation enhances motor skill acquisition over multiple days through an effect on consolidation. Proc Natl Acad Sci U S A, 106(5), 1590–1595. doi:10.1073/pnas.0805413106

Rezaee, Z., & Dutta, A. (2018). Transcranial Direct Current Stimulation of the Leg Motor Area - is it partly somatosensory? Annu Int Conf IEEE Eng Med Biol Soc, 2018, 4764–4767. doi:10.1109/EMBC.2018.8513195

Robertson, E. M. (2007). The serial reaction time task: implicit motor skill learning? J Neurosci, 27(38), 10073–10075. doi:10.1523/JNEUROSCI.2747-07.2007

Robertson, E. M., Pascual-Leone, A., & Miall, R. C. (2004). Current concepts in procedural consolidation. Nat Rev Neurosci, 5(7), 576–582. doi:10.1038/nrn1426

Robertson, E. M., Tormos, J. M., Maeda, F., & Pascual-Leone, A. (2001). The role of the dorsolateral prefrontal cortex during sequence learning is specific for spatial information. Cereb Cortex, 11(7), 628–635. doi:10.1093/cercor/11.7.628

Savic, B., & Meier, B. (2016). How Transcranial Direct Current Stimulation Can Modulate Implicit Motor Sequence Learning and Consolidation: A Brief Review. Front Hum Neurosci, 10, 26. doi:10.3389/fnhum.2016.00026

Seidel, O., & Ragert, P. (2019). Effects of Transcranial Direct Current Stimulation of Primary Motor Cortex on Reaction Time and Tapping Performance: A Comparison Between Athletes and Non-athletes. Front Hum Neurosci, 13, 103. doi:10.3389/fnhum.2019.00103

Stagg, C. J., Bachtiar, V., & Johansen-Berg, H. (2011). The role of GABA in human motor learning. Curr Biol, 21(6), 480–484. doi:10.1016/j.cub.2011.01.069

Taubert, M., Mehnert, J., Pleger, B., & Villringer, A. (2016). Rapid and specific gray matter changes in M1 induced by balance training. Neuroimage, 133, 399–407. doi:10.1016/j.neuroimage.2016.03.017

Teo, F., Hoy, K. E., Daskalakis, Z. J., & Fitzgerald, P. B. (2011). Investigating the Role of Current Strength in tDCS Modulation of Working Memory Performance in Healthy Controls. Front Psychiatry, 2, 45. doi:10.3389/fpsyt.2011.00045

Thielscher, A., Antunes, A., & Saturnino, G. B. (2015). Field modeling for transcranial magnetic stimulation: A useful tool to understand the physiological effects of TMS? Annu Int Conf IEEE Eng Med Biol Soc, 2015, 222–225. doi:10.1109/EMBC.2015.7318340

Yamamoto, S., Ishii, D., Ishibashi, K., & Kohno, Y. (2022). Transcranial Direct Current Stimulation of the Dorsolateral Prefrontal Cortex Modulates Cognitive Function Related to Motor Execution During Sequential Task: A Randomized Control Study. Front Hum Neurosci, 16, 890963. doi:10.3389/fnhum.2022.890963

Zhu, F. F., Yeung, A. Y., Poolton, J. M., Lee, T. M., Leung, G. K., & Masters, R. S. (2015). Cathodal Transcranial Direct Current Stimulation Over Left Dorsolateral Prefrontal Cortex Area Promotes Implicit Motor Learning in a Golf Putting Task. Brain Stimul, 8(4), 784–786. doi:10.1016/j.brs.2015.02.005

